# Gene2role: a role-based gene embedding method for comparative analysis of signed gene regulatory networks

**DOI:** 10.1101/2024.05.18.594807

**Authors:** Xin Zeng, Shu Liu, Bowen Liu, Weihang Zhang, Wanzhe Xu, Fujio Toriumi, Kenta Nakai

## Abstract

**Motivation:** Understanding the dynamics of gene regulatory networks (GRNs) across various cellular states is crucial for deciphering the underlying mechanisms governing cell behavior and functionality. However, current comparative analytical methods, which often focus on simple topological information such as the degree of genes, are limited in their ability to fully capture the similarities and differences among the complex GRNs.

**Results:** We present Gene2role, a gene embedding approach that leverages multi-hop topological information from genes within signed GRNs. Initially, we demonstrated the effectiveness of Gene2role in capturing the intricate topological nuances of genes using GRNs inferred from four distinct data sources. Then, applying Gene2role to integrated GRNs allowed us to identify genes with significant topological changes across cell types or states, offering a fresh perspective beyond traditional differential gene expression analysis. Additionally, we quantified the stability of gene modules between two cellular states by measuring the changes in the gene embeddings within these modules. In conclusion, our method augments the existing toolkit for probing the dynamic regulatory landscape, thereby opening new avenues for understanding gene behavior and interaction patterns across cellular transitions.

## 1. Introduction

Gene expression, a meticulously precise and intricately regulated process, is pivotal in maintaining cellular identity and facilitating differentiation. Within this complex regulatory framework lies a network of gene-to-gene interactions, known as the gene regulatory network (GRN). This network comprises nodes, representing genes, and edges, denoting the regulatory relationships between these genes. The nature of these regulatory relationships, whether activation or inhibition, is indicated by the sign of the edges.

GRNs can be constructed using a variety of methods based on diverse data sources (Emmert-Streib *et al*., 2014). First, GRNs can be built by collecting validated gene regulatory information from the literature. Due to its reliance on manually curated and experimentally verified data, this approach tends to produce relatively small networks, thereby limiting our exploration of the comprehensive regulatory relationship inside cells. Single-cell RNA sequencing (scRNA-seq) advancements have facilitated the reconstruction of GRNs encompassing thousands of genes by identifying gene co-expression patterns across various conditions or tissues (Akers and Murali, 2021). For instance, EEISP (Nakajima *et al*., 2021) constructs GRNs from scRNA-seq data based on co-dependency and mutual exclusivity of gene expression. However, this co-expression-based method struggles to differentiate direct from indirect gene regulatory relationships, complicating the accurate depiction of cellular dynamics. More recently, progress in single-cell multi-omics technology has further enhanced our ability to construct GRNs (Badia-i-Mompel *et al*., 2023). For example, CellOracle (Kamimoto *et al*., 2023) integrates scATAC-seq and scRNA-seq data, leveraging transcription factor (TF) binding motifs and co-expression information to infer GRNs, providing a more detailed and comprehensive understanding of gene regulatory mechanisms.

Downstream analysis of Gene Regulatory Networks (GRNs) is pivotal in uncovering gene functions. Prevalent analytical approaches emphasizing the topology of cell type-specific GRNs to identify key transcription factors (Aibar *et al*., 2017; Zeng *et al*., 2023) and gene modules (Langfelder and Horvath, 2008; Lemoine *et al*., 2021). However, these typical approaches, which focus on analyzing single network, often overlook the comparative analysis between GRNs across different cell states or types, thereby missing critical insights into the dynamics of regulatory mechanisms. Although methods exist for comparing GRNs between cell states (Duren *et al*., 2021; Kamimoto *et al*., 2023), they often focus solely on the direct topological information of genes, overlooking deeper structural connections (e.g., 1-hop and 2-hop neighbors), resulting in a shallow understanding of the complexity inherent in GRNs. To overcome this challenge, graph embedding techniques, which consider multi-hop connectivity, have been developed to project genes into an embedding space. These approaches preserve a richer representation of the original GRN information, thereby enabling a more precise quantification of gene distances.

However, current graph embedding approaches for GRNs rely on proximity principles (Wang *et al*., 2020; Wu *et al*., 2023; Gao *et al*., 2024), hindering the projection of genes from separate networks into closely positioned spaces for comparative analysis. Role-based network embedding methods such as struc2vec (Ribeiro *et al*., 2017) and SignedS2V (Liu *et al*., 2023) have introduced an advanced perspective by constructing a multi-layer weighted graph that reflects structural similarities among nodes at various depths. These methods facilitate embedding diverse networks into a unified space, allowing for nuanced comparisons of topological similarities across networks. Applying such advanced methodologies to compare GRNs among different cell states or types could significantly enhance our understanding of GRN dynamics.

In this article, we introduce Gene2role, a gene embedding method for signed GRNs, employing the frameworks from SignedS2V (Liu *et al*., 2023) and struc2vec (Ribeiro *et al*., 2017). We conducted experiments on GRNs generated from simulated network, manually curated networks, single-cell co-expression networks and single-cell multi-omics networks. These experiments demonstrated the ability of Gene2role to capture the topological information of GRNs. Additionally, we used Gene2role embeddings to analyze genes that exhibit structural variations across multiple GRNs and assessed the stability of gene modules across different cell states.

## 2. Materials and methods

The overall conceptual framework, shown in Fig. 1, consists of three major parts: network construction, embedding generation, and downstream analysis. Firstly, we introduce the network construction process from four data sources in Section 2.1. Then, we introduce the embedding generation framework in Sections 2.2-2.3, which applies SignedS2V to represent the topological features of genes in the signed GRN and to calculate the similarity between genes. Next, we explain the details of the gene embedding procedure in Sections 2.4-2.6 that adopts the struc2vec framework. The hyperparameters for experiments in our study are shown in Section 2.7. We showcase the downstream analysis based on gene embeddings at the gene and module levels, as depicted in Sections 2.8 and 2.9, respectively. Finally, we provide information about baseline methods and evaluation metrics in Sections 2.10 and 2.11.

**Figure 1.**
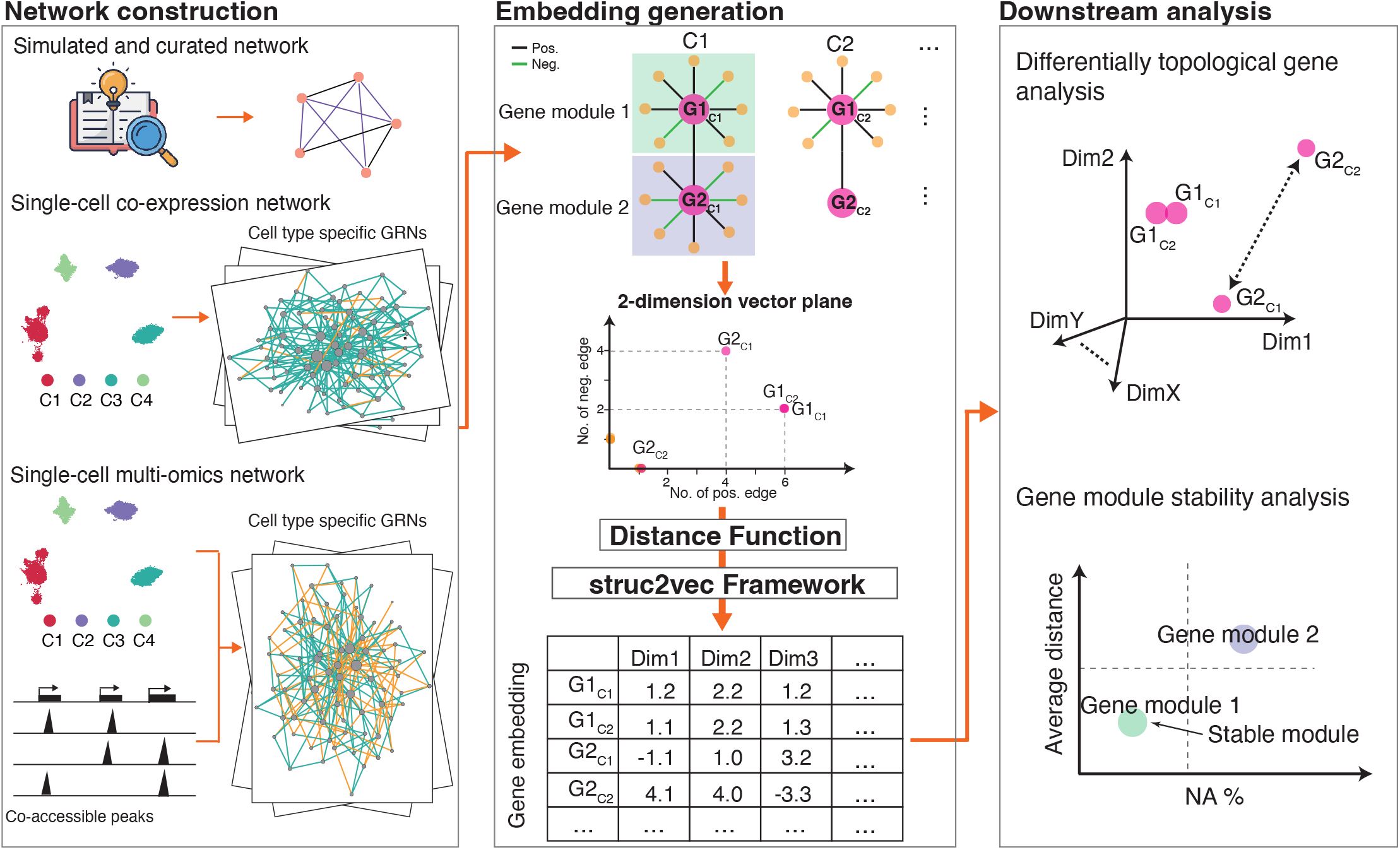
Overview of the Gene2role framework. Gene2Role is a multi-scale analysis framework for GRNs using a role-based graph embedding approach, which includes three components: network construction (left), embedding generation (middle), and downstream analysis (right). In network construction, one or more GRNs inferred by various methods are built, such as single-cell co-expression networks and single-cell multi-omics networks. In embedding generation, first, each gene is mapped to a 2-dimensional vector degree representing the number of positive and negative links. Next, the similarity between genes is evaluated using a series of distance functions that integrate multi-hop local topology information around the genes. Finally, the embedding is learned using the struc2vec framework. In downstream analysis, gene-level and gene-module-level analyses are performed. In gene-level analysis, differentially topological genes (such as G2) are extracted by comparing the distances of embeddings between GRNs. In gene-module-level analysis, the stability analysis of gene modules is performed by comparing the average distance between two GRNs and the proportion of genes that exist only in one of the GRNs (NA%) for pre-extracted gene modules. GRN: Gene regulatory network.

### 2.1 Network preparation

We constructed a simple simulated network to mimic the scale-free characteristics of GRNs, which comprising 31 genes. The four curated networks—hematopoietic stem cell (HSC) (Krumsiek *et al*., 2011), mammalian cortical area development (mCAD) (Giacomantonio and Goodhill, 2010), (ventral spinal cord) VSC (Lovrics *et al*., 2014), and (gonadal sex determination) GSD (Ríos *et al*., 2015)—containing between 5 to 19 genes, were downloaded from BELINE (Pratapa *et al*., 2020).

For single-cell RNA-seq data, we utilized the count matrix and cell type annotation data from two previous studies. Specifically, the dataset for human glioblastoma, as reported by (Nakajima *et al*., 2021), was collected at two distinct stages: the glioblastoma stem-like cells (0-hour) stage and the serum-induced differentiated (12-hour) stage. The datasets for human bone marrow mononuclear cells (BMMC) and human peripheral blood mononuclear cells (PBMC) were collected from (Granja *et al*., 2019). In the BMMC dataset, only Granulocyte-Macrophage Progenitors (GMPs) and CD14+ monocytes were kept, while in the PBMC dataset, all 10 cell types were retained. For each cell type, count matrices were generated using the 2000 highly variable genes. Subsequently, cell type-specific GRNs were constructed utilizing the EEISP (Nakajima *et al*., 2021) and Spearman correlation (Supplementary Note 1).

Single-cell multi-omics networks were obtained from CellOrcle (Kamimoto *et al*., 2023), which inferred them by integrating scRNA-seq data (Paul *et al*., 2015) and sci-ATAC-seq data (Cusanovich *et al*., 2018) derived from differentiating mouse myeloid progenitors. In brief, this dataset encompasses the differentiation of myeloid progenitors across 24 cell states, primarily highlighting the process of megakaryocyte and erythroid progenitors (MEPs) differentiating into erythrocytes, as well as GMPs differentiating into granulocytes. Within these networks, only connections exhibiting a p-value less than 0.01 were considered. Moreover, a selection criterion was applied to maintain only the top 2000 edges, chosen based on their highest absolute coefficient values. Consequently, these networks were composed of between 521 to 642 genes. The sign of edges was established based on the positive or negative values of their coefficient.

The detailed information on the networks utilized in the experiments described in this paper can be found in (Supplementary Table 1).

### 2.2 Gene topological representation in signed gene regulatory network (GRN)

Given a signed GRN represented as G = (*V, E*^+^, *E*^-^), where V = {*v*_1_,*v*_2_,···,*v*_*n*_ } denotes the set of genes. The sets E^+^ = {e |e ∈ *V* × *V*} and E^-^ = {e |e ∈ *V*× *V*} represent the positive and negative interactions between genes, respectively. To capture the topological nuances of each gene within the GRN, we introduce the concept of the signed-degree *d*, which is a 2-dimensional vector defined as:

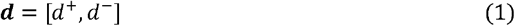

where *d*^+^and *d*^-^ are the positive and negative degrees, respectively. By adopting the signed-degree *d*, we map each gene from the signed GRNs to a point on the plane.

### 2.3 Gene topological similarity calculation

To quantify the topological similarity between genes within a GRN, we introduced the

distance function named Exponential Biased Euclidean Distance (EBED), which evaluates the zero-hop distance (*D*_0_) between the signed-degrees of two genes, 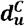 and 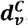, as follows:

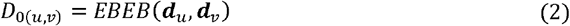

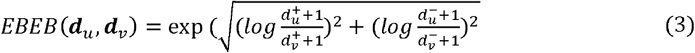

The rationale behind using EBED stems from the observation that GRNs are scale-free networks, and the degrees of their genes often follow a power-law distribution (Barabási and Oltvai, 2004). The EBED function initially employs a logarithmic transformation of the degrees to mitigate the effects of this distribution, then computes the Euclidean distance. An exponential function is subsequently applied to counterbalance the log transformation, thereby preserving the original proportionality of distances.

The topological identity of a gene is influenced not only by its direct connections but also by the broader topology that includes multi-hop neighborhoods. We define *R*_*k*_(*u*) as the sorted sequence of degrees for genes that are *k* hops away from gene *u*. Since the length of sorted sequence can be different for two genes, we employ dynamic time warping (DTW) (Salvador and Chan, 2007) to calculate the *k* -hop distance (*D*_*k*_, *k* > 0) between gene *u* and *v* using EBED as distance function:

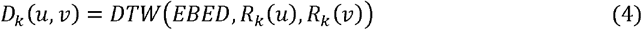

Then, the *k*-hop topological similarity between gene *u* and *v* can be calculated using the recursive formulation:

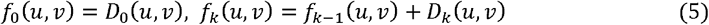

In this context, a lower value of *f*_*k*_(*u, v*) indicates a higher degree of similarity in the topology between gene *u* and *v*.

### 2.4 Multilayer graph construction

Next, we construct a multilayer weighted graph that encodes the topological information between genes. Each layer contains |*V*| genes, where the weight *w*_*k*_(*u, v*) for a link in the *k*-th layer (*k* > 0) between gene *u* and *v* is computed as:

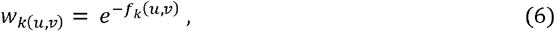

where a smaller value of *f*_*k*_ (*u, v*) indicates a higher topological similarity between gene *u* and *v*, thereby resulting in a greater weight for their link. For inter-layer connections, only the same genes are connected through directed links, with the weights of these links defined by:

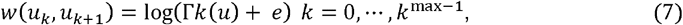

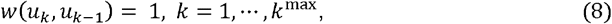

where *w*(*uk*,*uk*+l) corresponds to the weight between *uk* and *uk*+1, and *<u>w</u>*(*uk*,*uk*-l) corresponds to the weight between *uk* and *uk*-l. Γ*k*(*u*) denotes the number of links from gene *u* with weight exceeding the average link weight in the k-th layer. A high number of genes within the *k*-th layer share topological similarity with gene *u* leads to an increased Γ*k*(*u*), subsequently increasing the probability for searching topologically similar genes in the deeper layer. By setting *w*(*uk*,*uk*+l) to be greater than *w*(*uk*,uk-l), we encourage the exploration path in random walks to delve deeper rather than merely skimming the surface, aiming to uncover a wide range of topological relations between genes.

### 2.5 Sequence generation by random walk

Context sequences are generated using random walk on the weighted multilayer graph. The transition probability from gene *u* to *v* on layer *k*, denoted as *p*_*k*_(*u, v*), is determined by dividing *w*_*k*_(*u, v*) by the sum of the weights of all links connected to u within the layer:

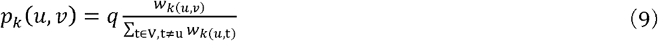

Here, *q* represents the intra-layer transition probability, set at 0.8. The inter-layer transition probability is set to 1-*q*, with the probabilities for upward and downward transitions being defined respectively as:

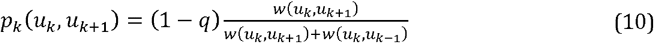

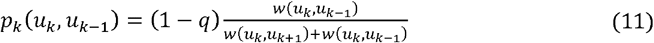

The transition probabilities ensure that every possible path originating from gene *u* is considered, with their probabilities summing to 1, reflecting the comprehensive exploration of connections both within and across layers.

### 2.6 Embedding learning

The Skip-Gram model (Mikolov *et al*., 2013) is finally used to create gene embeddings from the generated context sequences. The fundamental principle of this model is that the meaning of a word is determined by the words located closer to it in sentences. By leveraging this model, we can learn embeddings that encapsulate rich topological information, as context sequences consist of genes that are topologically similar. Importantly, this results in genes with similar topological structures being projected into closely situated points in the embedding space.

### 2.7 Hyperparameter for GRNs embedding experiments

For each experiment, we adjusted the parameters based on its complexity and the number of integrated networks (Supplementary Note 2).

### 2.8 Identification of differentially topological genes (DTGs)

Gene2role emphasizes the topological information of genes, allowing for genes that are distant or even unconnected from each other in a GRN to be positioned closely in the embedding space. To analyze the embedding of the same gene *u* from *C* cell types, we first determine the center of embeddings denoted as ***cen***_***u***_. For each cell type, we calculate the Euclidean distance 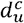 from the embedding 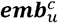 of gene *u* to the ***cen***_***u***_:

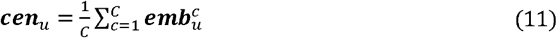

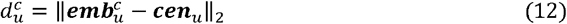

We then determine the average distance 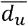, to provide a summary measure of the distance of gene *u* embeddings from the centroid across all cell types.

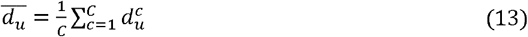

For C=2, genes with the largest distances ranking in the top 10% are defined as DTGs. For C>2, we further calculate the standard deviation *σ*_*u*_ to quantify the topological variability of gene *u*:

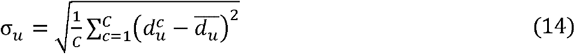

Genes located in the top 5th percentile of average distances are classified as DTGs. Additionally, genes whose standard deviation exceeds a certain empirical threshold are also identified as DTGs. The high average distances indicate that these genes consistently exhibit significant differences across all cell states, suggesting a universal role change. Conversely, a high standard deviation points to substantial variability in specific cell states, implying that these genes may only undergo significant role changes under certain conditions, while remaining relatively stable in others.

When comparing DTGs with differentially expressed genes (DEGs), we identified DEGs using Seurat (Hao *et al*., 2021) pipeline (Supplementary Note 3).

### 2.9 Gene module stability analysis

To quantify the stability of gene modules between two cell types or states, we initially define an anchor cell type (*c*_1_). Then, we use the Louvain algorithm, a proximity-based clustering method, to organize the GRN of the anchor cell type into gene modules;

Within these modules, genes typically collaborate in certain specific functions. (Lemoine *et al*., 2021). Gene modules containing fewer than 10 genes will be excluded. For each remaining gene module *m*, we calculate the average distance 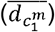 of genes between the two cell types using the following equation:

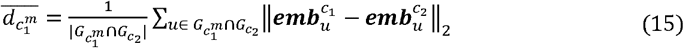

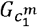 is the gene set for module *m* in cell type *c*_1_, and 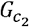 is the whole gene set in cell type *c*_2_. |·| represents the size of set.

Moreover, the absence of gene embeddings in cell types other than the anchor may indicate significant changes in their roles. To quantify these changes, we define 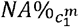 as the percentage of genes within a gene module *m* that only occur in the anchor cell type *c*_1_.

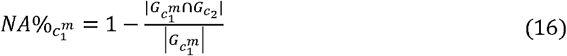

We combine the average gene distance and NA% as a measure to assess the overall change in roles of gene modules that were initially clustered together in anchor cell type when observed in another cell type.

To assess the biological functions of a gene module, we employ the *compareCluster* function from the clusterProfiler package (Yu *et al*., 2012). This analysis employed the *enrichGO* function, which targets biological processes (BPs) using org.Hs.eg.db and org.Mm.eg.db as the gene ontology database for dataset from human and mouse, respectively. We apply the Benjamini-Hochberg method to adjust p-values, with a q-value cutoff set at 0.05 to determine significance. The top five significant Gene Ontology (GO) terms for each gene module were retained for further analysis and visualization.

### 2.10 Baseline methods

We compared Gene2role to two graph embedding methods: struc2vec(Ribeiro *et al*., 2017) and BESIDE (Chen *et al*., 2018). In brief, struc2vec is a role-based embedding method that does not consider sign information, whereas BESIDE considers the sign information and is based on proximity embedding.

### 2.11 Evaluation metrics

The topological features of a gene in a signed GRN were evaluated using using the following metrics: degree centrality (+), degree centrality (-), betweenness centrality, eigenvector centrality, degree assortativity (+), degree assortativity (-), clustering coefficient (+), clustering coefficient (-) (Supplementary Note 4).

## 3 Results

### 3.1 Gene2role captures the topological information of GRNs

To verify that Gene2role accurately captures the topological information of genes within GRNs, we analyzed GRNs derived from four distinct data sources: 1. one simulated network, and 2. four curated networks based on experimentally validated interactions (Section 3.1.1), 3. co-expression networks generated from single-cell RNA sequencing data (Section 3.1.2), 4. multi-omics networks derived from multi-omics data (Section 3.1.3).

#### 3.1.1 Simulated and curated GRN

To provide a clear overview of our network embedding study, we selected one simulated network (Fig. 2A) and four simple curated networks (Fig. 2C, Supplementary Fig. 1A, C, E) for analysis. We compared Gene2role to two other methods, struc2vec (Ribeiro *et al*., 2017) and BESIDE (Chen *et al*., 2018), by setting the embedding dimension to 2 for all networks (Fig.2B, D, Supplementary Fig. 1B, D, F). Overall, Gene2role effectively positioned genes with similar connectivity patterns—marked by both positive and negative edges—within proximity in the embedding space. In contrast, the disregard of struc2vec for edge sign failed to accurately capture the topological nuances of genes. BESIDE, despite considering edge signs, its proximity-based embedding strategies, which bring genes with positive connections closer together while pushing those with negative connections farther apart, failed to capture the topological information of genes. For example, in simulated network, genes S9 and S15 were closely positioned by Gene2role due to their single negative edge. Gene S0 was projected near genes S6-S8 and S10-S14, reflecting their equal distance to negative edges. Conversely, although struc2vec closely aligned genes with similar positive topological information, it overlooked the S9 and S15 due to its exclusive focus on positive connections. BESIDE placed S9 far from S1 and S15 far from S2 due to their respective negative connections, clearly demonstrating how the method handles negative links between genes. Additionally, in the HSC network, despite their spatial separation, Gene2role clustered Fli1, Eklf, cJun, and EgrNab together due to their shared configuration of one negative and one or two positive edges. Struc2vec, however, placed cJun and EgrNab, each with a single positive edge, distantly from Fli1 and Eklf, who both had two positive edges, indicating a disregard for edge sign complexity. In contrast, although BESIDE grouped Fli1 and Eklf together, it also mixed in other genes, demonstrating its inability to effectively capture topological information. Furthermore, in the GSD network, Gene2role positioned DHH and PGD2 adjacent to each other because of their only positive linkage to a single gene and without any negative links. Struc2vec similarly grouped FGF9 with DHH and PGD2 solely based on their single positive edge, overlooking the negative edge. Conversely, BESIDE, by considering network proximity and edge signs, positioned AMH and FGF9 near PGD2 and DHH.

**Figure 2.**
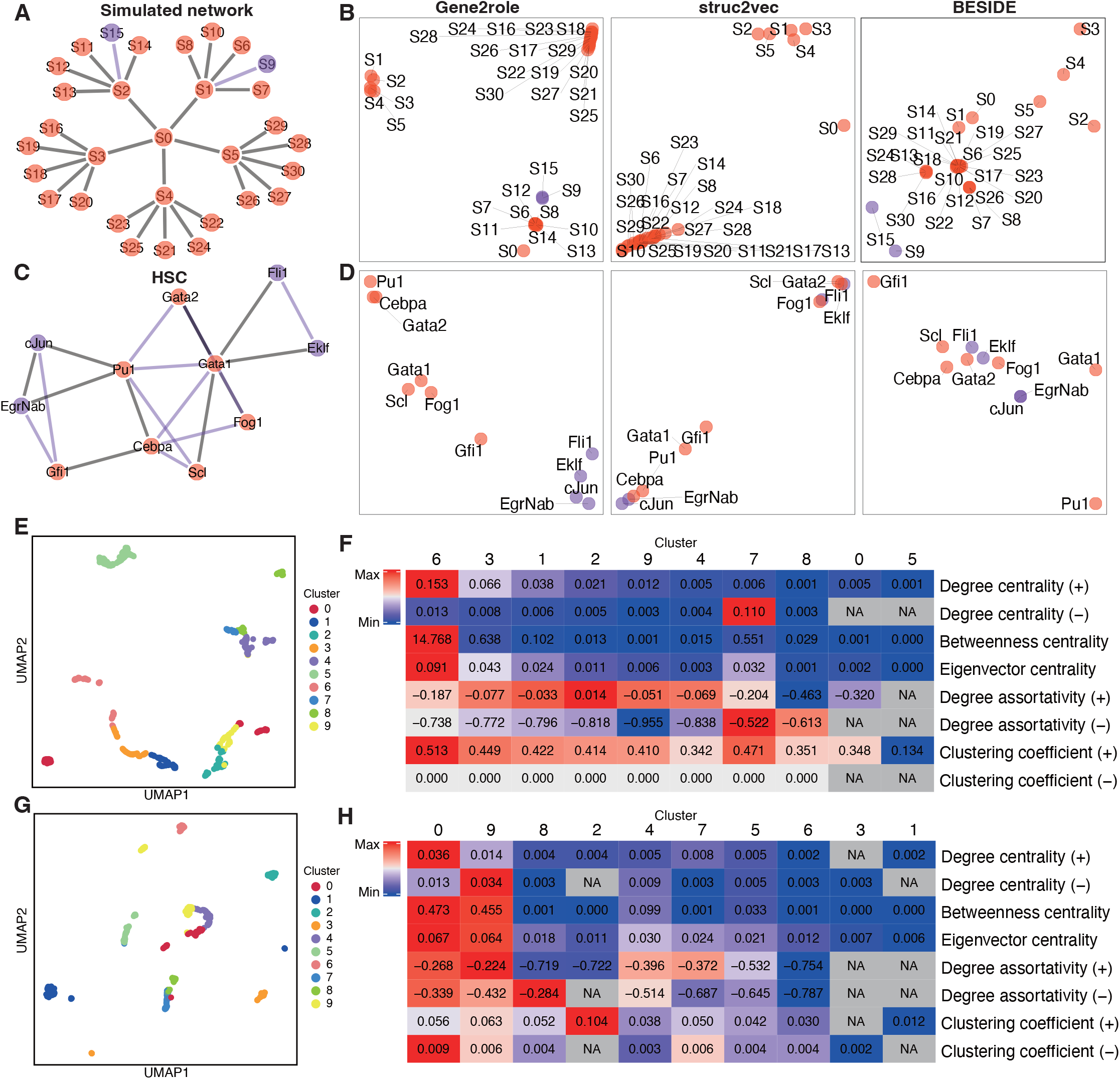
Analysis of GRN embeddings from networks derived through various methods. **A, C** Simulated tree GRN (**A**) and HSC network(**C**). **B, D** 2D embeddings for networks from (**A**) and (**C**), respectively. The embeddings were generated by Gene2role, struc2vec, and BESIDE, respectively. **E, G** K-means (K=10) clustering of embeddings from B cells in human PBMC dataset (**E**) and Ery_0 stage in multi-omics dataset (**G**) displayed in UMAP. **F, H** Heatmap displaying the average values of 8 network feature metrics for the 10 clusters of genes from (**E**) and (**G**) within the GRN. The color scaling within each row is determined by the maximum and minimum values of that row. UMAP: Uniform manifold approximation and projection

#### 3.1.2 Single-cell co-expression network

Having demonstrated the effectiveness of Gene2role in simple networks, we extended our exploration to more complex GRNs derived from single-cell RNA-seq data. We first applied Gene2role to a GRN that was inferred from B cells in the human PBMC dataset using EEISP. Gene embeddings were organized into ten distinct clusters using the K-means algorithm (Fig. 2E). We then calculated the average value of eight topological metrics for the genes within each cluster, reflecting their interconnectedness within the original GRN. The heatmap revealed unique metric enrichment across the clusters (Fig. 2F), highlighting the distinct topological attributes inherent to each cluster. Notably, genes in cluster 6 exhibited significant prominence in positive degree centrality, indicating that they were central genes with a high density of positive interactions. In contrast, cluster 7 was distinguished by its considerable negative degree centrality and minimal positive connections, suggesting a group of genes primarily linked by negative edges. Cluster 2, enriched with high positive degree assortativity but minimal negative degree centrality, predominantly comprised genes positioned on the outer edges with positive links. Furthermore, cluster 9, characterized by the lowest negative degree assortativity and minimal negative degree centrality, was identified as a peripheral cluster primarily connected through negative relationships.

To test the performance of our method across different GRN inference methods, we applied Gene2role to the GRN that was inferred from B cells in the human PBMC dataset using Spearman correlation. The gene distribution across clusters demonstrated a consistent topological structure, mirroring the patterns discerned in the EEISP results (Supplementary Fig. 1G, H). For instance, the characteristics of cluster 7 were similar to those of cluster 6 in the EEISP-derived clusters, delineating a hub of genes with densely packed positive linkages. Similarly, cluster 5 corresponded to the previously identified cluster 7, characterized by a richness in negative interactions. Additionally, cluster 0 shared traits with cluster 2, encompassing genes predominantly linked through positive edges and situated at the periphery of the GRN. Finally, the genes within cluster 1 resembled those in cluster 9, forming a peripheral group predominantly characterized by negative edges.

#### 3.1.3 Single-cell multi-omics network

We further explored more sophisticated GRNs that was inferred by integrating single-cell RNA-seq and single-cell ATAC-seq data. We used Gene2role to analyze the Ery_0 stage of the multi-omics GRNs, which was collected from CellOracle. We segmented genes into ten groups using K-means and calculated the average value of eight key metrics for each cluster (Fig. 2G, H). Genes in cluster 0 functioned as a hub predominantly associated with positive edges, whereas genes in cluster 9 functioned as their negative-edge counterpart. Cluster 8, characterized by high negative degree assortativity and low centrality, represented a group of genes on the periphery of the negative network. Conversely, cluster 2, lacking negative connections and possessing the highest positive clustering coefficient, comprised a cohesive group of genes with similar functions and low degrees of connectivity.

### 3.2 Identification of DTGs between paired states using Gene2role embeddings

Given the ability of Gene2role to group genes by topological patterns, we merged GRNs from two cell types to explore the gene role changes across these networks. We applied Gene2role to the human glioblastoma dataset, which comprised GRNs from 0-hour and 12-hour stages. We computed the pairwise distances for each gene and extracted the DTGs by the top 10% largest distances (Fig. 3A). 66 DTGs were identified and 50 of them were overlapped with DEGs (Fig. 3B). For instance, there was a significant drop in CD164 expression and its network connections at 12-hour stage (Supplementary Fig. 2A, B), which aligns with its known role in glioblastoma proliferation (Wang *et al*., 2019). Additionally, we observed that although 16 genes exhibited minor expression differences between the two cell types, their connection patterns within GRNs underwent significant changes. Specifically, the expression levels of DKK3 remained consistent between both cell types (Fig. 3C); however, its network connectivity diminished at the 12-hour stage (Fig 3D). Moreover, we identified that 397 genes that underwent significant changes in expression levels, yet their topological structures within the GRNs remained unchanged. For instance, while the expression level of EGR1 decreased, its network connectivity was maintained (Supplementary Fig. 2C, D).

**Figure 3.**
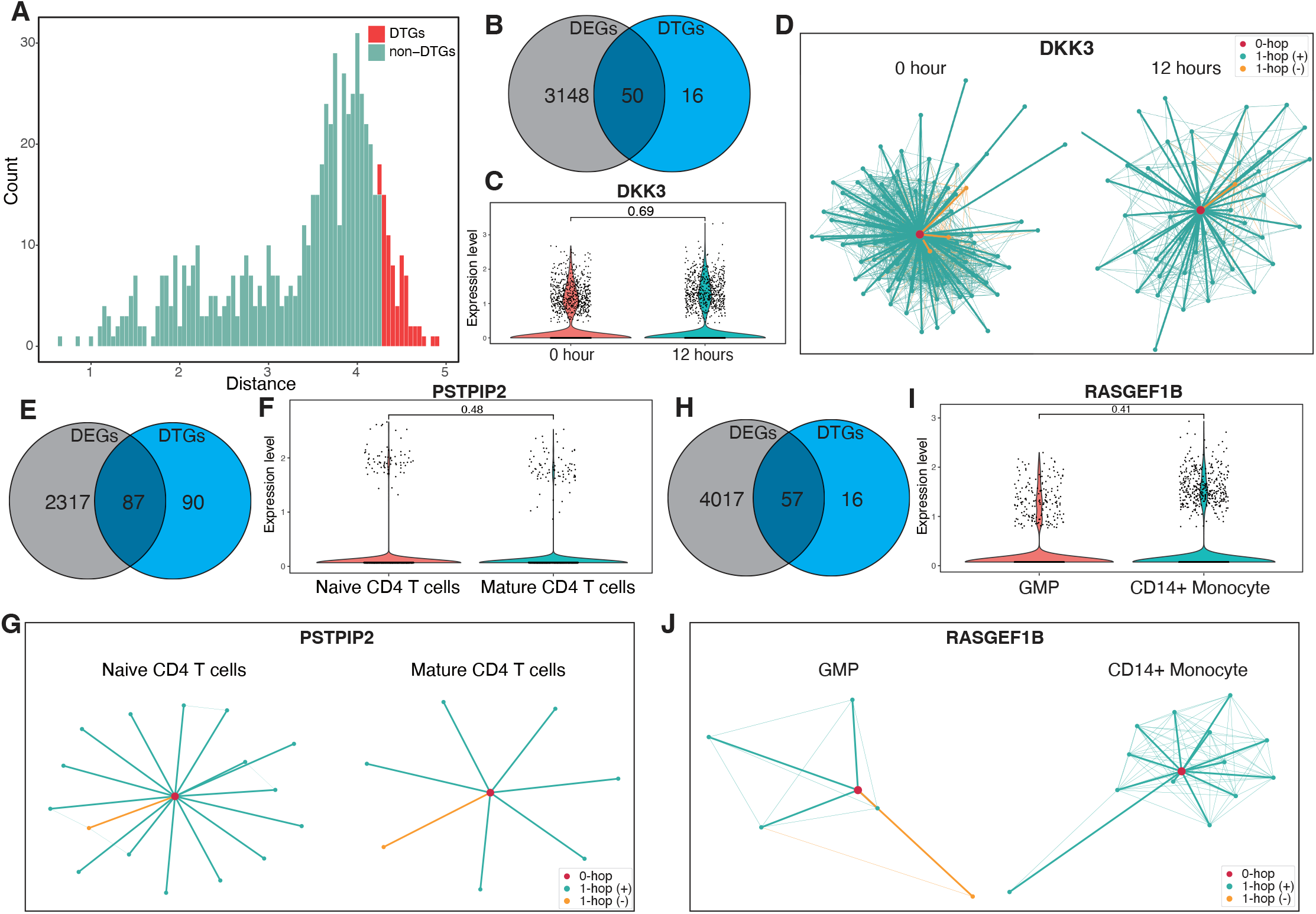
Distinctive analysis of DTGs in comparison with DEGs across two cell types. **A** histogram of the frequency distribution of Gene2role embedding pair distances in GRNs at 0-hour and 12-hour stage within a human glioblastoma dataset, with the top 10th percentile of average distances designated as DTGs. **B, E**, and **H** Venn diagrams depict the intersection between DEGs and DTGs identified in human glioblastoma at 0-hour and 12-hour stage (**B**), in human PBMCs comparing naïve CD4 T cells with mature CD4 T cells (**E**), and in human BMMC comparing GMPs with CD14+ Monocytes (**H**), respectively. **C, F**, and **I** Examples of genes that are exclusively DTGs not overlapping with DEGs from the intersecting sets in (**B**), (**E**), and (**H)**, respectively. **D, G, J** 1-hop network structures corresponding to the DTG shown in **C, F**, and **I** DTGs: differentially topological genes; DEGs: differentially express genes; PBMC: peripheral blood mononuclear cells; BMMC: human bone marrow mononuclear cells; GMPs: Granulocyte-macrophage progenitors

We extended our application of Gene2role to analyze the topological shifts of genes between naïve CD4 T cells and mature CD4 T cells within the human PBMC dataset. We discovered that 87 DTGs overlapped with DEGs, indicating a robust pattern across cell types (Fig 3E). For example, CARD16 showcased an increase in both expression level and connectivity when comparing naïve to mature CD4 T cells (Supplementary Fig. 2E, F). Furthermore, despite 90 genes showing only minor expression differences between these two cell types, significant alterations in their network connectivity were evident. A case in point is PSTPIP2, which maintained consistent expression levels across both cell types (Fig 3F), yet its network connectivity notably decreased in mature CD4 T cells (Fig 3G). In addition, an analysis revealed that 716 genes experienced significant shifts in expression levels without corresponding changes in their topological structures within the GRNs. SP140, for instance, exhibited an increased expression level, while its network position remained stable (Supplementary Fig. 2G, H).

In the analysis of GMPs and CD14+ Monocytes from the human BMMC dataset, we identified 57 DTGs that overlapped with DEGs (Fig 3H), including genes such as ETS1 that displayed significant alterations in both expression levels and topological structures (Supplementary Fig. 2I, J). Conversely, 16 DTGs, which were not DEGs, such as RAGEF1B, showed significant topological changes without a corresponding shift in expression levels (Fig 3I, J). Moreover, within the DEGs, 524 genes, exemplified by RETN, underwent substantial changes in expression while their topological configurations were preserved (Supplementary Fig. 2K, L).

### 3.3 Identification of DTGs among cell types

Having analyzed gene role changes between two cell types, we extended our study to merged GRNs from multiple cell types. We applied Gene2role to the integrated GRNs from the human PBMC dataset, which comprise 10 cell types. Initially, we identified 33 DTGs by focusing on the top 5th percentile of average distances (Fig. 4A). These genes exhibited significant variability across the 10 cell types. For instance, the gene FGFBP2 had a higher count of negative edges in NK cells and central memory CD8 T cells compared to other cell types, while it showed the fewest positive edges in B cells (Supplementary Fig. 3A). Moreover, analysis revealed that seven genes exhibited a

**Figure 4.**
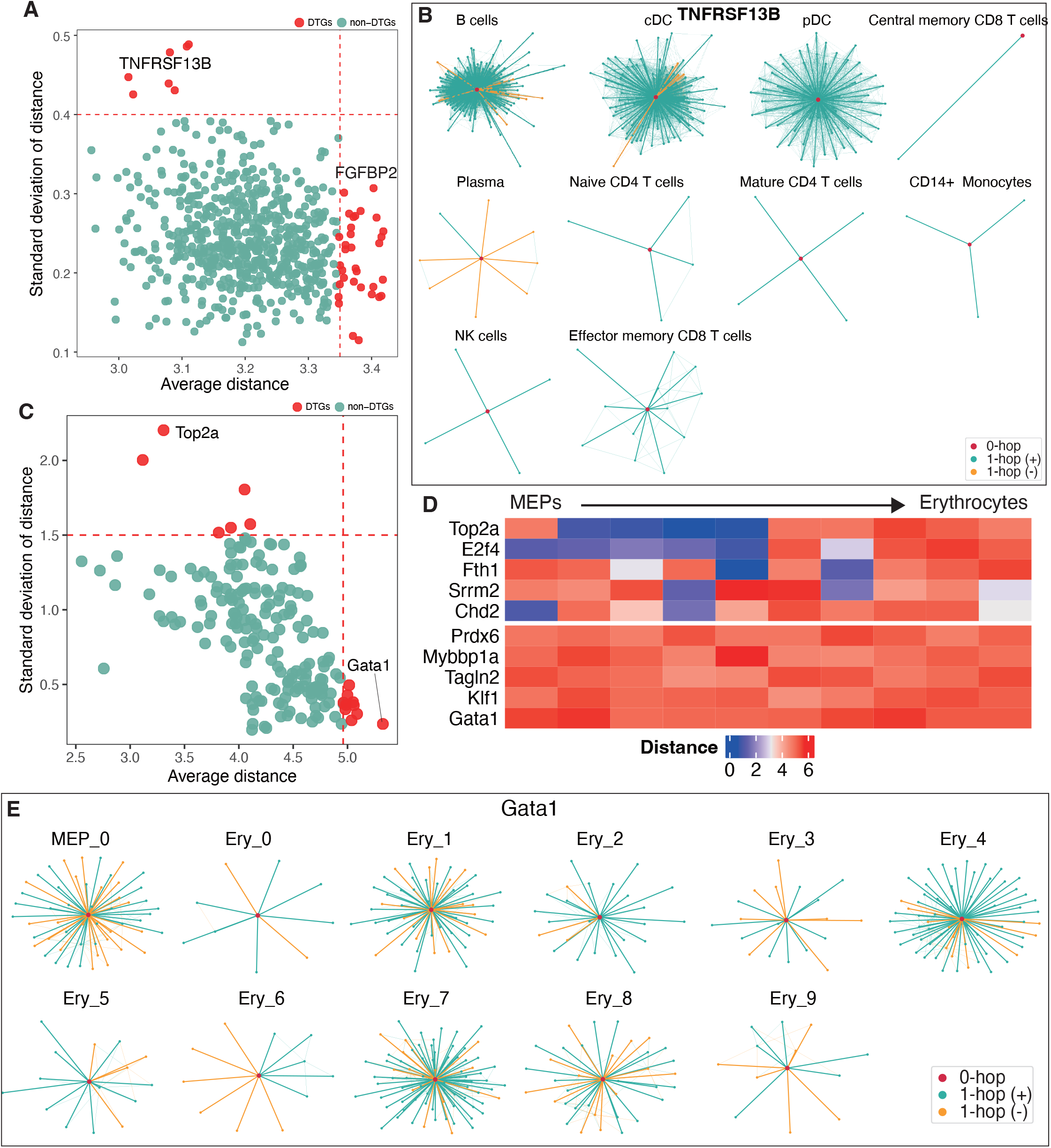
Comparative analysis of gene distance across multiple cell types. **A** Scatter plot of the average and standard deviation of gene distances within human PBMC dataset across 10 cell types, with genes in the top 5th percentile of average distances or standard deviation greater than 0.4 were categorized as DTGs. **B** 1-hop network structures of TNFRSF13B in ten GRNs from human PBMC datase. **C** Scatter plot of the average and standard deviations of gene distances between each pair of adjacent cell types in the differentiation trajectory of MEPs. **D** Heatmap of two patterns of gene role changes during MEP differentiation: the top heatmap represents genes with high variance and low average distance, while the bottom heatmap depicts genes with low variance but high average distance. **E** 1-hop network structures of TNFRSF13B in eleven GRNs from multi-omics dataset MEPs: Megakaryocyte-erythroid progenitors

standard deviation in topological distance exceeding 0.4, yet an average distance below 3.1, indicative of substantial structural dispersion among these cell types. For example, TNFRSF13B functioned as a hub gene in B cells, conventional dendritic cells (cDCs), and plasmacytoid dendritic cells (pDCs), but maintained fewer connections in other cell types (Fig. 4B).

Subsequently, we used Gene2role embeddings to investigate significant topological changes in genes during the differentiation of MEPs and GMPs in the multi-omics dataset. For each gene, we calculated the average and standard deviation of distance between adjacent cellular state across 11 sequential stages during erythrocyte differentiation. By focusing on the top 5th percentile of genes in terms of average distance or gene with high standard deviation of distance, we identified two distinct patterns of DTGs (Fig. 4C, D). The first pattern, characterized by a high standard deviation in distance but a relatively small average distance, suggests that the roles of these genes may remain unchanged during certain developmental stages, only to shift abruptly thereafter. For instance, Top2a exhibited no significant topological changes from the Ery_0 to Ery_4 stages but underwent noticeable changes post-Ery_5 (Supplementary Fig. 3B), suggesting a role shift after this stage. The second pattern is characterized by consistently large changes in role distance throughout development, indicating dynamic and substantial fluctuations in gene roles. For example, the gene Gata1 exhibited significant periodic fluctuations in the number of its connections (Fig. 4E). Previous studies, such as (Fujiwara *et al*., 1996), have shown that Gata1 promotes the differentiation of erythroid cells, and our findings corroborate its dynamic involvement in this process .

Similarly, we calculated the average distance and standard deviation of each gene between seven adjacent cellular states during granulocyte differentiation. We observed topological shift patterns similar to those identified in erythrocyte differentiation (Supplementary Fig. 4A, B). For example, structural variations in gene Smarcc1 were primarily observed before the Gran_0 stage and stabilized between Gran_0 and Gran_2, suggesting a temporal shift in its topological significance (Supplementary Fig. 4C). Additionally, Eif3g exhibited continuous changes throughout its development, reinforcing its active role in the differentiation process (Supplementary Fig. 4D).

### 3.4 Evaluation of gene module stability

After analyzing topological variations of the same gene across different GRNs, we focused on the overall topological changes in a single gene module between two GRNs. In the human glioblastoma dataset, we designated the 0-hour stage as the anchor cell type, clustered the GRN into seven gene modules, and calculated the average pairwise gene embedding distance and NA% between the 0-hour and 12-hour stages. Each gene module, characterized by distinct NA% and average distances, suggests shifts in the roles of these gene modules during differentiation (Fig. 5A). For example, gene module 5 exhibited a relatively large mean distance and high NA%, suggesting substantial changes in the topological structure of the genes within this module. Gene Ontology (GO) analysis for gene module 5 highlighted metabolic processes, notably cholesterol and sterol biosynthesis (Fig. 5B). Consequently, we infer that the changes in the topological structure of gene module 5 may indicate reduced in sterol biosynthesis capabilities within the cells at the 12-hour stage. Previous research has demonstrated that patient-derived glioblastoma stem cells (GSCs) activate cholesterol biosynthesis more extensively than differentiated glioblastoma cells (Gu *et al*., 2023). Conversely, gene module 0 exhibited a lower mean distance and NA%, indicating minimal changes in the topological structure of the genes within this module. GO analysis revealed that gene module 0 was enriched in fundamental cellular processes, including chromosome segregation and nuclear division (Fig. 5B). This observation suggests that the functions of chromosome segregation and nuclear division are relatively stable between the 0-hour and 12-hour stages. Additionally, when we designated the 12-hour stage as the anchor cell type, the clustering resulted in five main gene modules. We observed that gene module 1 was relatively stable between the 0-hour and 12-hour stages (Supplementary Fig. 5A), exhibiting similar biological processes to those of gene module 0 when 0-hour served as the anchor cell type (Supplementary Fig. 5B). This observation suggests that they may represent the same gene module, consistently maintaining a stable topological structure and biological function throughout the differentiation process. In contrast, gene module 9, characterized by its enrichment in biological functions related to extracellular matrix and structure organization, was relatively unstable.

**Figure 5.**
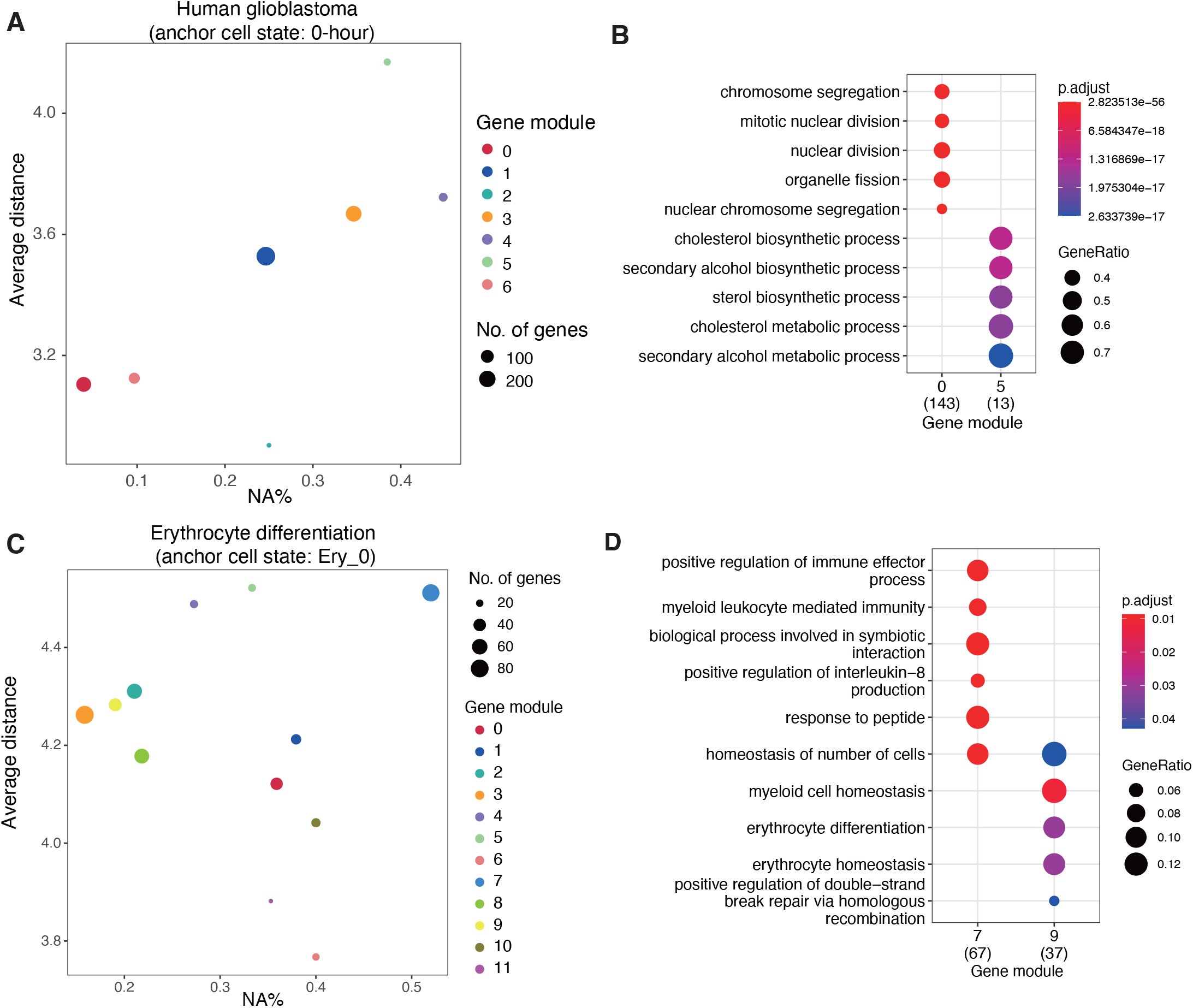
Gene module stability analysis using Gene2role embeddings. **A** Scatter plot depicting the mean distance and the percentage of genes exclusively found in the anchor cell type (NA%) for seven gene modules in a human glioblastoma dataset, using the 0-hour stage as the anchor cell type and comparing against the 12-hour stage. **B** Dot plot of the top 5 significant biological processes from the GO analysis for gene modules 0 and 5 in **A. C** Scatter plot for twelve gene modules during MEP differentiation, with the Ery_0 stage as the anchor cell type and comparing against the Ery_9 stage, illustrating the mean distance and percent of genes unique to the anchor cell type (NA%). **D** Dot plot for the top 5 significant biological processes from the GO analysis of gene modules 7 and 9 in **C**.

To explore gene module stability during erythrocyte differentiation, we established Ery_0 stage as the anchor cell type and identified 12 gene modules. We calculated the average distances and NA% for Ery_3 (Supplementary Fig. 5C), Ery_6 (Supplementary Fig. 5D), and Ery_9 (Fig. 5C) in relation to the Ery_0 stage across the 11 gene modules. We observed that several gene modules, including gene module 9, maintained stable positions throughout the development process. GO analysis indicated that gene module 9 was enriched in biological process related to erythrocyte differentiation homeostasis, suggesting a continuous role in driving erythrocyte differentiation (Fig. 5D). In contrast, gene module 7 undergoes significant role changes and involved in biological processes related to immune regulation, suggesting that its immunomodulatory functions may be diminished during differentiation.

By setting GMP_0 as the anchor cell type and computing the average distances and NA% from Gran_3 stage, we further explored the stability of 11 gene modules during granulocytes differentiation (Supplementary Fig. 5E). Similarly, we observed that gene module 2, which was enriched in biological process related to mononuclear cell differentiation, was unstable during granulocytes differentiation (Supplementary Fig. 5F). This observation suggests that role of genes driving mononuclear cell differentiation has changed during granulocytes differentiation. Gene module 6, which was stable and enriched in fundamental metabolic processes, implies that this module provides essential metabolites during differentiation.

## Discussion

In this study, we introduced Gene2role, a gene embedding method that utilizes the topological attributes of genes within gene regulatory networks (GRNs). Our findings demonstrate that Gene2role effectively captures the topological information within GRNs constructed from four distinct data sources. By facilitating the integration of GRNs across various cell states and types, Gene2role enables robust comparative analysis at two levels: first, it identifies genes exhibiting significant topological discrepancies among cell types; secondly, it evaluates the stability of gene modules across two cellular states.

Our analysis of DTGs between two cell types across multiple datasets consistently revealed three gene patterns: those exhibiting significant changes in both expression and topology, those with altered expression but stable topology, and those with stable expression but significant topological changes. These observations suggest a relative independence between changes in gene roles within cellular networks and alterations in their expression level. Importantly, our approach identifies genes that exhibit notable shifts in their topological roles within the co-expression network, despite showing no significant changes in expression levels. Such genes may undergo functional changes that traditional differential gene expression analyses frequently overlook. Thus, our method provides a valuable complementary perspective by focusing on topological shifts within GRNs. One limitation of ourmethod is that it identifies only those genes with structural variations that maintain at least one connection in the networks. Consequently, it may miss changes in genes that are embedded in some cell types but absent in others. Future research could focus on inductively quantifying distances for genes selectively connected across different cell types to enhance our comprehension of how gene role changes across cellular contexts.

We examined topological shifts within gene modules to assess their stability across two cellular states. Combined with Gene Ontology (GO) analysis, this stability assessment enables a nuanced understanding of the relationship between the functional dynamics of gene modules and shifts in cellular states. For instance, in analyzing erythrocyte differentiation from the multi-omics dataset, we observed that gene module 9 maintained a low average distance and NA% between Ery_0 and Ery_9, and it was implicated in the promotion of erythrocyte differentiation by GO analysis. (Fig. 5D) Hence, this gene module may play a stable role in driving erythrocyte differentiation. In our stability analysis of a gene module between two cellular states, we focused on the proximity structures with the anchor cell type. However, a deeper investigation into the proximity information in the non-anchor cell type could reveal proximity-based shifts between cell types. By integratively investigating the topological and proximity shifts between cell types, we can further discern the coherence or dysfunction of gene modules, thus providing specific insights into their stability.

Although our method has been tested in GRNs inferred from single-cell RNA-seq and multi-omics datasets, it is equally applicable to spatial transcriptomics data (Larsson *et al*., 2021). Recent analyses of spatial transcriptomics have concentrated on inferring spatial domains by integrating expression information, spatial information, and histological information(Long *et al*., 2023; Yang *et al*., 2023). These spatial domains can serve as a basis for identifying spatial GRNs as inputs for our method, thereby enabling a deeper understanding of topological variations among GRNs in spatial contexts.

## Acknowledgement

Computational resources were provided by the supercomputer system SHIROKANE at the Human Genome Center, Institute of Medical Science, the University of Tokyo.

## Data availability

The edgelists of curated networks were downloaded from BEELINE (Pratapa *et al*., 2020). For single-cell RNA-seq data, the count matrix and metadata of human glioblastoma, human PBMC, and human BMMC were collected from GEO databaset (GSE144623, GSE139369). The edgelists generated from multi-omics data were collected from CellOrcle (Kamimoto *et al*., 2023). Codes generated for this project is available on GitHub (https://github.com/liushu2019/g2r_work).

