## Supplemental Information for "Gene2role: a role-based gene embedding method for comparative analysis of signed gene regulatory networks"

### Supplementary Notes

#### Supplementary Note 1: Details of data preparation for single-cell co-expression network inference

***Generation of count matrix***. We mainly analyzed three single-cell RNA-seq datasets, originating from human glioblastoma (Nakajima *et al.*, 2021), human bone marrow mononuclear cells (BMMC), and human peripheral blood mononuclear cells (PBMC) (Granja *et al.*, 2019). For the human glioblastoma dataset, a total of 4311 cells were downloaded, including 2102 glioblastoma stem cells collected at 0 hours and 2209 serum-induced differentiated glioblastoma cells collected at 12 hours. The BMMC dataset included 1323 Granulocyte-Macrophage Progenitors (GMP) and 2681 CD14+ monocytes. The PBMC dataset contained 9436 cells across 10 cell types, specifically including 56 pDCs, 189 cDCs, 1050 B cells, 50 plasma cells, 2321 mature CD4 T cells, 2371 naive CD4 T cells, 747 effector memory CD8 T cells, 1336 central memory CD8 T cells, 1793 NK cells, and 3432 CD14+ monocytes.

For the analysis of integrated network embeddings of two cell types, three datasets were utilized: human glioblastoma; GMP and CD14+ monocytes from the BMMC dataset; and naive CD4 T cells and mature CD4 T cells from the PBMC dataset. Following log normalization and scaling, top 2000 highly variable genes between the paired cell types were identified using *FindVariableFeatures* from Seurat (Hao *et al.*, 2021). Count matrices were subsetted to a dimension of No. cells × 2000 high variable genes for each cell type under study.

For the analysis of integrated network embeddings more than two cell types, only the PBMC dataset was used. Using the *FindVariableFeatures* function, after data were log-normalized and scaled, the top 2000 highly variable genes across all cell types were identified. Count matrices were subsetted to a dimension of No. cells × 2000 high variable genes for each cell type under study.

***Inference of co-expression networks.*** Our network inference approach and parameters were tailored based on the number of networks integrated downstream. For single network embedding, we employed EEISP (Nakajima *et al.*, 2021) and Spearman correlation to analyze the count matrix of B cells from the human PBMC dataset. Specifically, in the Spearman correlation analysis, gene pairs within certain thresholds were selected to construct the network, utilizing the top 1% for positive edges and the top 0.1% for negative edges. A similar methodology was adopted for EEISP, where gene pairs were selected based on the top 1% of the co-dependency index (CDI) for positive edges and the top 0.1% of the exclusively expressed index (EEI) for negative edges. For two networks embedding, we utilized EEISP, selecting gene pairs based on the top 1% of the CDI for positive edges and the top 0.1% of the EEI for negative edges. In multi-networks embedding, we adjusted the selection criteria to include the top 3% of gene pairs by CDI for positive edges, while still selecting the top 0.1% by EEI for negative edges.

#### Supplementary Note 2: Details in network embedding *hyperparameters*

***Hyperparameters***: In our study, we employed the signedS2V(Liu *et al.*, 2023) and struc2vec (Ribeiro *et al.*, 2017) frameworks for network embedding, which involve several hyperparameters that can significantly influence the effectiveness of the embeddings. These include '--until-layer', which defines the number of hops; '--num-walks', specifying the number of random walks initiated from each gene; '--walk-length', which sets the length of each random walk; '--window-size', determining the size of the window for the skip-gram model; and '--dimensions', which sets the dimensionality of the embeddings. '--non-scale-free' is the parameter to control the usage of EBED, which can be added if the input GRN is relatively small and is not a scale-free network.

Additionally, our model incorporates three optimizable options, further details of which can be found in the documentation for struc2vec.

***Simulated and curated networks.*** For all simulated and curated networks analyzed, the parameters were consistently set to --walk-length 80, --window-size 5, and --dimensions 2. Specifically, for simulated networks, '--until-layer' was adjusted to 5 and '--num-walks' to 20. In the case of curated networks, '--until-layer' was set to 2, with '--num-walks' increased to 100.

***Single-cell co-expression networks and single cell multi-omics networks.*** For the embedding of both single-cell co-expression networks and single-cell multi-omics networks, the hyperparameters were uniformly set as follows: --walk-length 80, --window-size 10, and --dimensions 128. Specifically, '--until-layer 3' was selected to determine the exploration depth within the network structure, while '--num-walks' was established at 100 to ensure comprehensive network traversal.

#### Supplementary Note 3: Identification of differentially expressed genes (DEGs)

We used Seurat (Hao *et al.*, 2021) to identify DEGs. Specifically, for the analysis of integrated network embeddings of two cell types, three datasets were normalized and scaled through *SCTransform* function. DEGs between two cell types were identified using *FindMarkers* with adjusted p-value of 0.05*.* For the analysis of integrated network embeddings of more than two cell types, human PBMC dataset was normalized and scaled through the *SCTransform* function. Marker genes were identified using the *FindAllMarkers* function, applying a threshold of an adjusted p-value < 0.05 and setting 'only.positive = TRUE' to focus exclusively on positive markers.

#### Supplementary Note 4: Evaluation metrics

We used 8 metrics to assess the topological information of a gene in the GRN.

***Degree centrality***: Degree centrality measures the importance or influence of a gene $u$within a GRN by quantifying the number of direct connections, or edges, it has. In a signed network, we distinguish between degree centrality (+) ${DC}_{u}^{+}$and degree centrality (-) ${DC}_{u}^{-}$for both signs:

$${DC}_{u}^{+}=\sum_{i}^{N} A_{ui}^{+}$$

$${DC}_{u}^{-}=\sum_{i}^{N} A_{ui}^{-}$$

where $A_{ui}^{+}$ and $A_{ui}^{-}$are the positive and negative adjacency matrix elements, respectively. And N is the total number of genes in the GRN.

***Betweenness centrality***. Betweenness centrality (Brandes, 2001) is a measure that identifies the frequency with which a gene appears on the shortest paths between pairs of other genes within a network. The betweenness centrality of a gene $u$ can be calculated as:

$${BC}_{u}=\sum_{s\neq u\neq t} \frac{\sigma_{st}(u)}{\sigma_{st}}$$

where $\sigma_{st}$ represents the total count of weighted shortest paths from gene $s$ to gene $t$, and $\sigma_{st}\left( u \right)$ counts how many of those paths pass through gene $u$. In this context, edge weights, derived from their signs, are utilized to determine the distances between gene pairs, leading the shortest path algorithm to navigate through a mix of positive and negative edges. Therefore, a high (low) value of positive (negative) betweenness centrality indicates the prevalent of a gene involvement in the shortest paths that predominantly include positive (negative) edges, highlighting its role as a pivotal connector across different network regions.

***Eigenvector centrality.*** Eigenvector centrality measures the influence of a gene based on the importance of its neighbors. A gene with high eigenvector centrality is connected to other genes with high eigenvector centrality.

$${EC}_{u}=\frac{1}{\lambda}\sum_{i}^{N} A_{ui}{EC}_{j}$$

where $\lambda$ is the largest eigenvalue of the adjacency matrix $A=A^{+}+A^{-}$.

***Degree assortativity.*** Degree assortativity (Newman, 2003) quantifies the propensity for genes within a network to link to others with similar degrees, essentially measuring the pattern of connectivity where similar genes are more likely to connect. In the context of signed networks, evaluating positive and negative degree assortativity separately becomes crucial because of the unique characteristics of each type of connection. This metric is relevant to the network as a whole; hence, to understand local structures better, we analyze the degree assortativity within the ego network of each gene. The calculation of Degree assortativity is expressed as:

$$r=\frac{1}{\sigma_{a}^{2}}\sum_{ij} ij(e_{ij}-a_{i}b_{j})$$

where $e_{ij}$ denotes the proportion of edges that link genes of degree $i$ with those of degree $j$, $a_{i}$(and $b_{j}$) represents the fraction of edges emanating from (or directed to) genes of degree $i$ (or $j$), $\sigma_{a}^{2}$is the variance of the distribution of $a_{i}$, calculated by:

$$\sigma_{a}^{2}= \sum_{i} i^{2}a_{i}-{(\sum_{i} ia_{i})}^{2}$$

***Clustering coefficient.*** The clustering coefficient (Schank and Wagner, 2005) is a metric that quantifies the prevalence of triangular connections surrounding a gene $u$, essentially gauging how interconnected a gene's neighbors are. Like Degree assortativity, we differentiate between positive and negative clustering coefficients to capture the nuances of connections in signed GRNs. The formula for the clustering coefficient of gene $u$is defined as:

$$c_{u}=\frac{No. of triangles connected to gene u}{No. of triples centered on gene u}$$

For any given gene $u$ with a degree of $k_{u}$, the denominator, representing the potential number of triples centered on gene $u$, is calculated as:

$$\left( \frac{k_{u}}{2} \right)=\frac{k_{u}(k_{u}-1)}{2}$$

### Supplementary Figures

#### Supplementary Figure 1

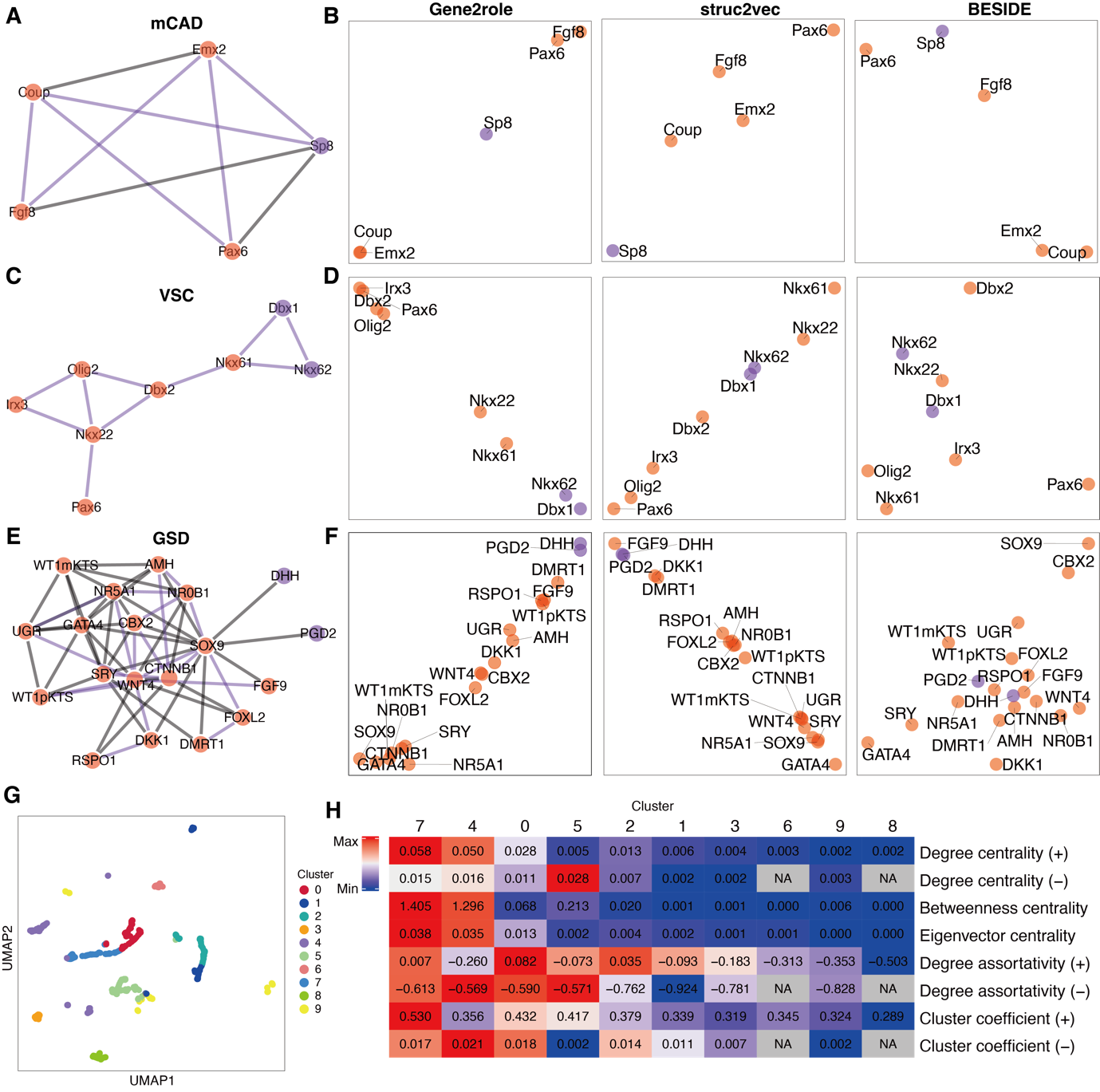

***Supplementary Fig. 1.*** ***A****,* ***C****,* ***E*** *mCAD network (****A****), VSC network (****C****), and GSD network(****E****).* ***B****,* ***D, F*** *2D embeddings for networks from (****A****), (****C****) and (****E****), respectively. The embeddings were generated by Gene2role, struc2vec, and BESIDE, respectively.* ***G*** *K-means (K=10) clustering of embeddings generated from spearman co-expression network from B cells in human PBMC dataset displayed in UMAP.* ***H*** *Heatmap displaying the average values of 8 network feature metrics for the 10 clusters of genes from (****G****) within the GRN. The color scaling within each row is determined by the maximum and minimum values of that row.*

*UMAP: uniform manifold approximation and projection*

#### Supplementary Figure 2

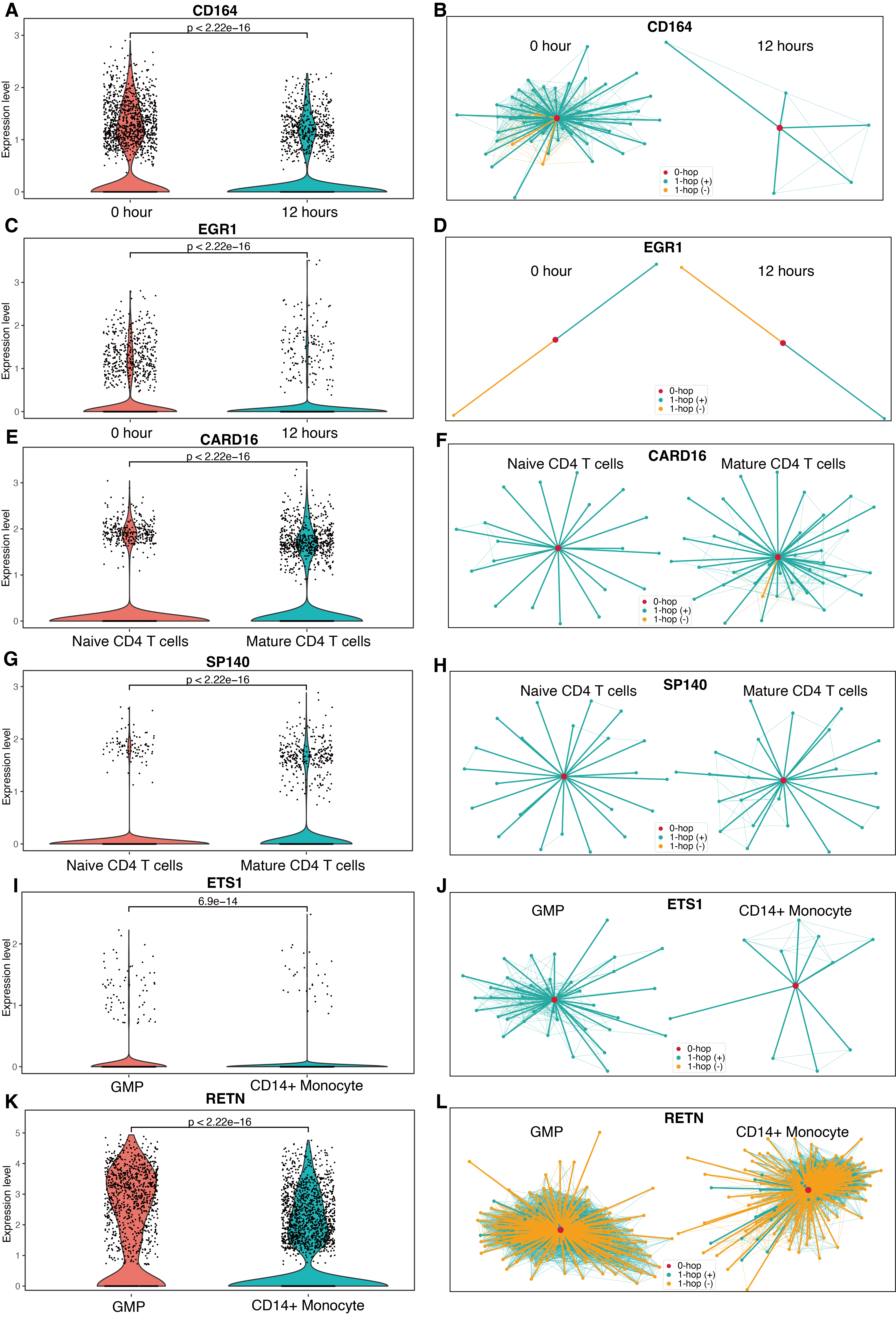

***Supplementary Fig. 2*** ***A****,* ***E****, and* ***I*** *Examples of genes identified as both DTGs and DEGs, highlighting the intersections of these sets in* ***Fig. 3B, E, H****, respectively.* ***B****,* ***F****,* ***J*** *1-hop network structures corresponding to the DTG shown in* ***A****,* ***E****, and* ***I****.* ***C, G, K*** *Examples of genes exclusively DEGs not overlapping with DTGs in human glioblastoma, human PBMC, human BMMC dataset.* ***D****,* ***H****,* ***L*** *1-hop network structures corresponding to the DTG shown in* ***C****,* ***G****, and* ***J****.*

#### Supplementary Figure 3

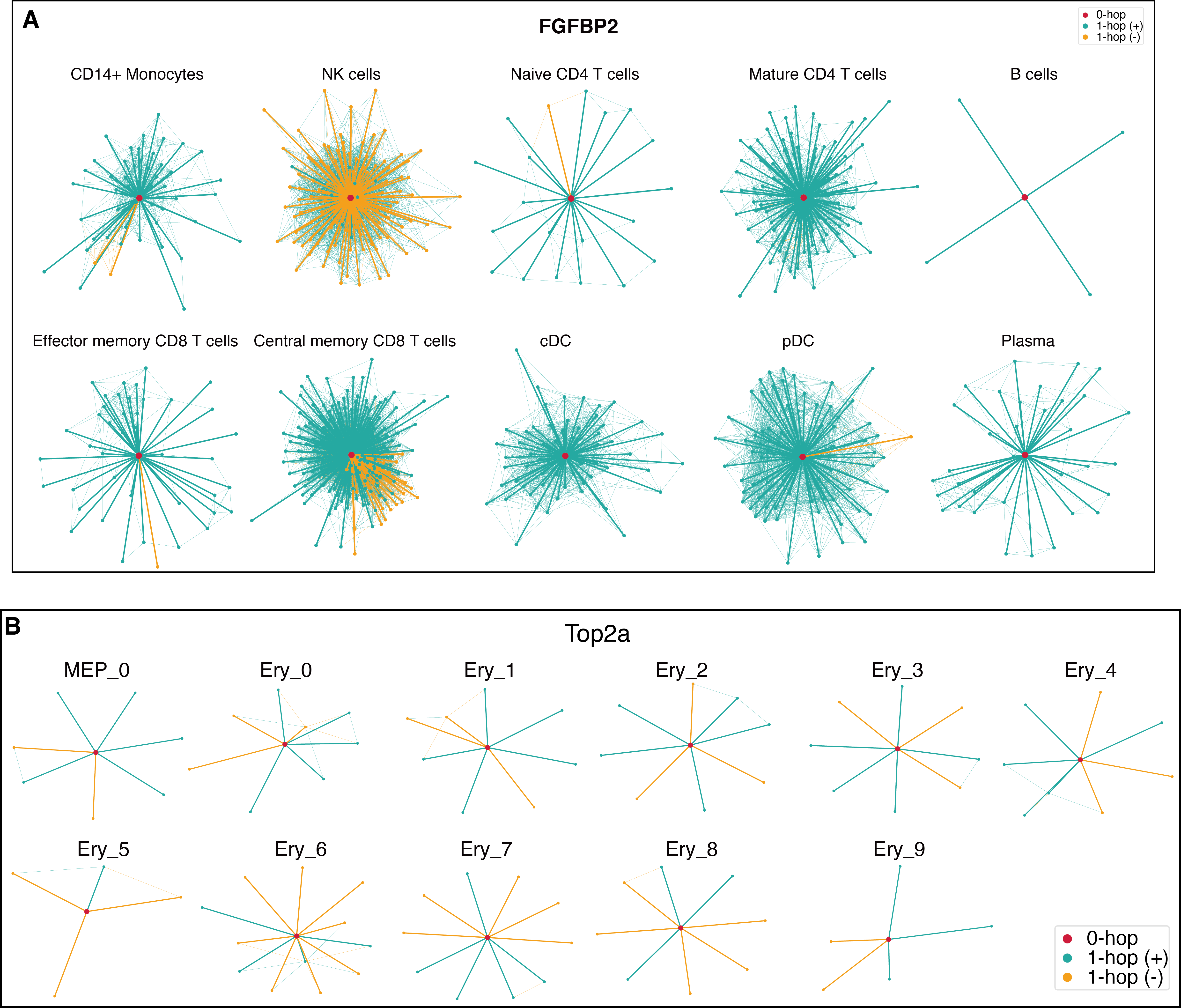

***Supplementary Fig. 3. A*** *1-hop network structures of FGFBP2 with high average distance across 10 different GRNs in human PBMC dataset.* ***B*** *1-hop network structures of Top2a with high standard deviation of distance across 11 different GRNs in multi-omics dataset.*

#### Supplementary Figure 4

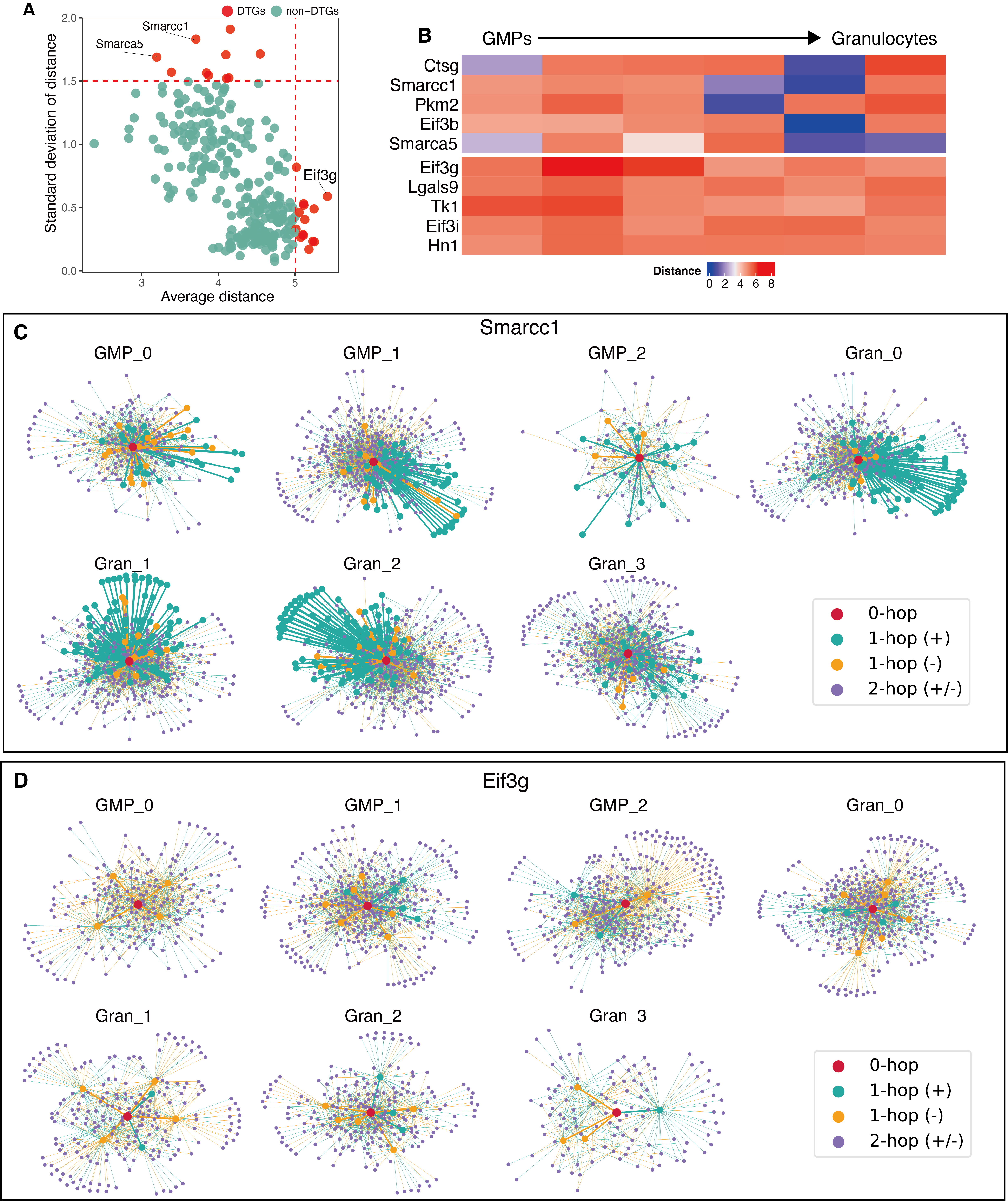

***Supplementary Fig. 4.*** ***A*** *Scatter plot of the average and standard deviations of gene distances between each pair of adjacent cell types in the differentiation trajectory of GMPs.* ***B*** *Heatmap of two patterns of gene role changes during GMP differentiation: the top heatmap represents genes with high variance and low average distance, while the bottom heatmap depicts genes with low variance but high average distance.* ***C*** *2-hop network structures of Smarcc1 with high average distance across 7 different GRNs in multi-omics dataset.* ***D*** *2-hop network structures of Eif3g with high standard deviation of the distance across 7 different GRNs in multi-omics dataset.*

*GMP: Granulocyte-Macrophage Progenitors*

#### Supplementary Figure 5

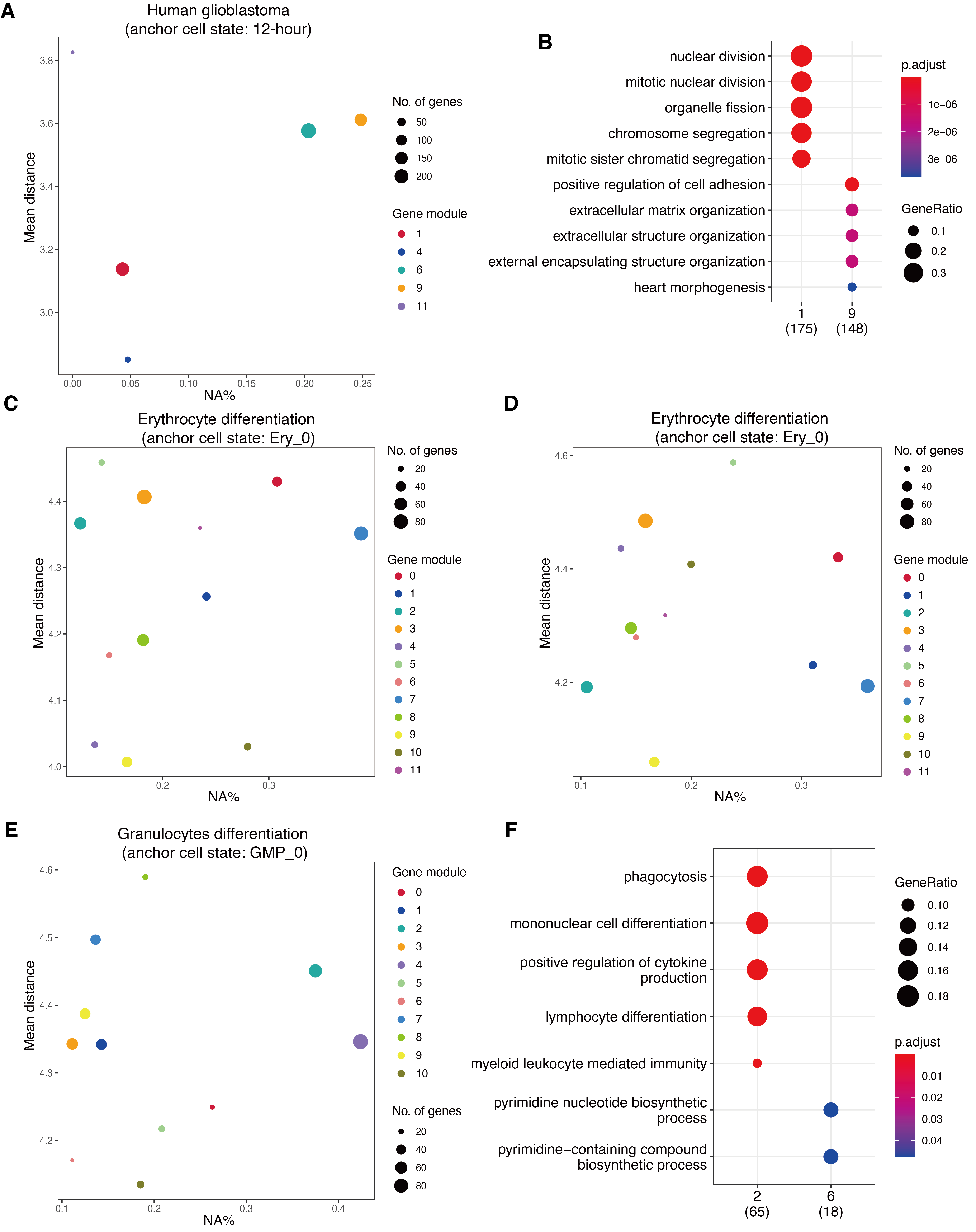

***Supplementary Fig. 5.*** ***A*** *Scatter plot depicting the mean distance and the percentage of genes exclusively found in the anchor cell type (NA%) for seven gene modules in a human glioblastoma dataset, using the 12-hour stage as the anchor cell type and comparing against the 0-hour stage.* ***B*** *Dot plot of the top 5 significant biological processes from the GO analysis for gene modules 1 and 9 in* ***A****.* ***C, D*** *Scatter plot for twelve gene modules during MEP differentiation, with the Ery_0 stage as the anchor cell type and comparing against the Ery_3 (****C****) and Ery_6 (****D****) stage, illustrating the mean distance and percent of genes unique to the anchor cell type (NA%)****. E*** *Scatter plot for twelve gene modules during GMP differentiation, with the GMP_0 stage as the anchor cell type and comparing against the Gran_3 stage, illustrating the mean distance and percent of genes unique to the anchor cell type (NA%).* ***F*** *Dot plot for the top 5 significant biological processes from the GO analysis of gene modules 2 and 6 in* ***E****.*

### Supplementary Tables

#### Supplementary Table 1: Statistics of GRNs

| **Experiment**  **type** | **Dataset** | **Cell types** | **# genes** | **# common genes** | **# edges** | **Pos. edges %** |
| --- | --- | --- | --- | --- | --- | --- |
| Single network | Simulated network | NA | 31 | NA | 30 | 93.3 |
|  | HSC | NA | 11 | NA | 21 | 47.6 |
|  | mCAD | NA | 5 | NA | 9 | 33.3 |
|  | VSC | NA | 8 | NA | 10 | 0 |
|  | GSD | NA | 19 | NA | 59 | 66.1 |
|  | Human PBMC | B cells  (EEISP) | 749 | NA | 18338 | 85.7 |
|  | Human PBMC | B cells  (Spearman) | 1199 | NA | 14621 | 91.0 |
|  | Multi-omics | Ery_0 | 552 | NA | 2000 | 57.7 |
| Two networks | Human glioblastoma | 0h,  12h | 884  906 | 655 | 31040  28458 | 87.9  86.8 |
|  | Human PBMC | N. CD4 T,  M. CD4 T | 1888  1861 | 1770 | 27858  35578 | 87.0  88.8 |
|  | Human BMMC | GMPs,  CD14 Mono. | 1122  1122 | 729 | 36092  27630 | 89.4  86.9 |
| More than two networks | Human PBMC | All | 1251-1676 | 658 | 34374-79938 | 93.3-95.6 |
|  | Multi-omics | All | 521-642 | 111 | 2000 | 51.3-64.2 |
